# Female gene networks are expressed in myofibroblast-like smooth muscle cells in vulnerable atherosclerotic plaques

**DOI:** 10.1101/2023.02.08.527690

**Authors:** Ernest Diez Benavente, Santosh Karnewar, Michele Buono, Eloi Mili, Robin J. G. Hartman, Daniek Kapteijn, Lotte Slenders, Mark Daniels, Redouane Aherrahrou, Tobias Reinberger, Barend M. Mol, Gert J. de Borst, Dominique P. V. de Kleijn, Koen H. M. Prange, Marie A. C. Depuydt, Menno P. J. de Winther, Johan Kuiper, Johan L. M. Björkegren, Jeanette Erdmann, Mete Civelek, Michal Mokry, Gary K Owens, Gerard Pasterkamp, Hester M. den Ruijter

**Affiliations:** Laboratory of Experimental Cardiology, University Medical Center Utrecht, Utrecht University, the Netherlands; Robert M. Berne Cardiovascular Research Center, University of Virginia, Charlottesville, VA, USA; Central Diagnostic Laboratory, University Medical Center Utrecht, Utrecht University, Utrecht, the Netherlands; Center for Public Health Genomics, University of Virginia, Charlottesville, VA, USA; Institute for Cardiogenetics, University of Lübeck, Lübeck, Germany; A.I. Virtanen Institute for Molecular Sciences, University of Eastern Finland, Finland; Department of Vascular Surgery, University Medical Centre Utrecht, Utrecht, Utrecht University, the Netherlands; Division of BioTherapeutics, Leiden Academic Centre for Drug Research, Leiden University’ Leiden, the Netherlands; Department of Genetics and Genomic Sciences, Icahn School of Medicine at Mount Sinai, New York, NY, USA; Department of Medicine, Karolinska Institutet, Karolinska Universitetssjukhuset, Huddinge, Sweden; Department of Biomedical Engineering, University of Virginia, Charlottesville, VA, USA

## Abstract

Women presenting with coronary artery disease (CAD) more often present with fibrous atherosclerotic plaques, which are currently understudied. Phenotypically modulated smooth muscle cells (SMCs) contribute to atherosclerosis in women. How these phenotypically modulated SMCs shape female versus male plaques is unknown. Here, we show sex-stratified gene regulatory networks (GRNs) from human carotid atherosclerotic tissue. Prioritization of these networks identified two main SMC GRNs in late-stage atherosclerosis. Single-cell RNA-sequencing mapped these GRNs to two SMC phenotypes: a phenotypically modulated myofibroblast-like SMC network and a contractile SMC network. The myofibroblast-like GRN was mostly expressed in plaques that were vulnerable in females. Finally, mice orthologs of the female myofibroblast-like genes showed retained expression in advanced plaques from female mice but were downregulated in male mice during atherosclerosis progression. Female atherosclerosis is driven by GRNs that promote a fibrous vulnerable plaque rich in myofibroblast-like SMCs.

---

Women are predominantly protected from cardiovascular disease at young ages. However, once in their 70s the incidence of coronary artery disease in women surpasses that of men^1^. This suggests a strong interaction of sex and age^2,3^, where menopause is a known inflection point ^4^.

Eroded plaques are most prevalent in women^5^ but also contribute substantially to acute coronary syndromes (ACS) in younger men^6,7^. These plaques are characterized by their fibrous composition and lack of substantial calcification^8^. Despite the importance of symptomatic fibrous plaques, the biological processes that lead to fibrous plaques and erosion have received less attention than those leading to plaque rupture^9^.

Previously, we demonstrated in aortic tissue of patients suffering from coronary artery disease the dominance of expression of female specific gene regulatory networks in smooth muscle cells (SMCs) and Endothelial cells (ECs)^10^. Endothelial to mesenchymal transition (EndMT) is considered a potential mechanism of plaque erosion^11^ and was found enriched in the female-specific gene regulatory networks (GRNs)^10^. Furthermore, key driver genes from the SMC main network were upregulated in female SMCs and expressed in phenotypically modulated SMCs in *Klf4* knock-out lineage-tracing mice^12^, suggesting that they play a role in SMCs plasticity and that this is an important mechanism in female plaques^10^.

Despite these recent advances, it is unknown how these GRNs contribute to the distinct female plaque phenotype. We hypothesize that the newly-identified female SMC network contributes to plaque phenotypic differences between sexes and informs on plaque vulnerability in women. In this study, we applied systems biology on human atherosclerotic plaques from the carotid artery and show differential SMC biology in females compared to males. We prioritized two female GRNs that are active in extracellular matrix (ECM)-producing myofibroblasts and involved in vulnerable plaque phenotypes. In line with these findings, we observed that key driver genes of the female GRNs remained expressed in plaques isolated from lineage-tracing female mice during atherosclerosis progression but were downregulated in plaques from male mice.

## Results

### Patient population and female GRNs in carotid atherosclerotic plaques

To create robust and reliable female-biased GRNs of atherosclerosis, we used a systems biology approach in an equally powered cohort of atherosclerotic plaque tissues from women and men^10^. We selected all women with carotid plaque RNAseq data (n=158) and age-matched these with 158 men from the Athero-Express carotid endarterectomy biobank^13,14^ (See Online methods, **Figure 1**). Differences in the male and female populations with respect to their risk factor profiles included significant higher high-density lipoprotein (HDL) in females (1.3 vs 1.0 mmol/L; *p*-value (*p*)<0.001), a higher proportion of current smoking in females (46% vs 32% in males; p<0.001) and higher prevalence of alcohol consumption in males (72% vs 48%; *p*<0.001) (**Table S1**). A higher prevalence of CAD history was found in males (compared to females (37.3% vs. 24.7%, *p*=0.021) as well as significantly higher use of anticoagulants in males compared to females (17% vs. 7%, p=0.007). As reported previously^8^, female plaques were more fibrous - highlighted by a high content of SMCs and low content of macrophages (MΦ). Conversely, male plaques showed a phenotype with higher prevalence of plaque haemorrhage and fat content (**Table S1**).

**Figure 1.**
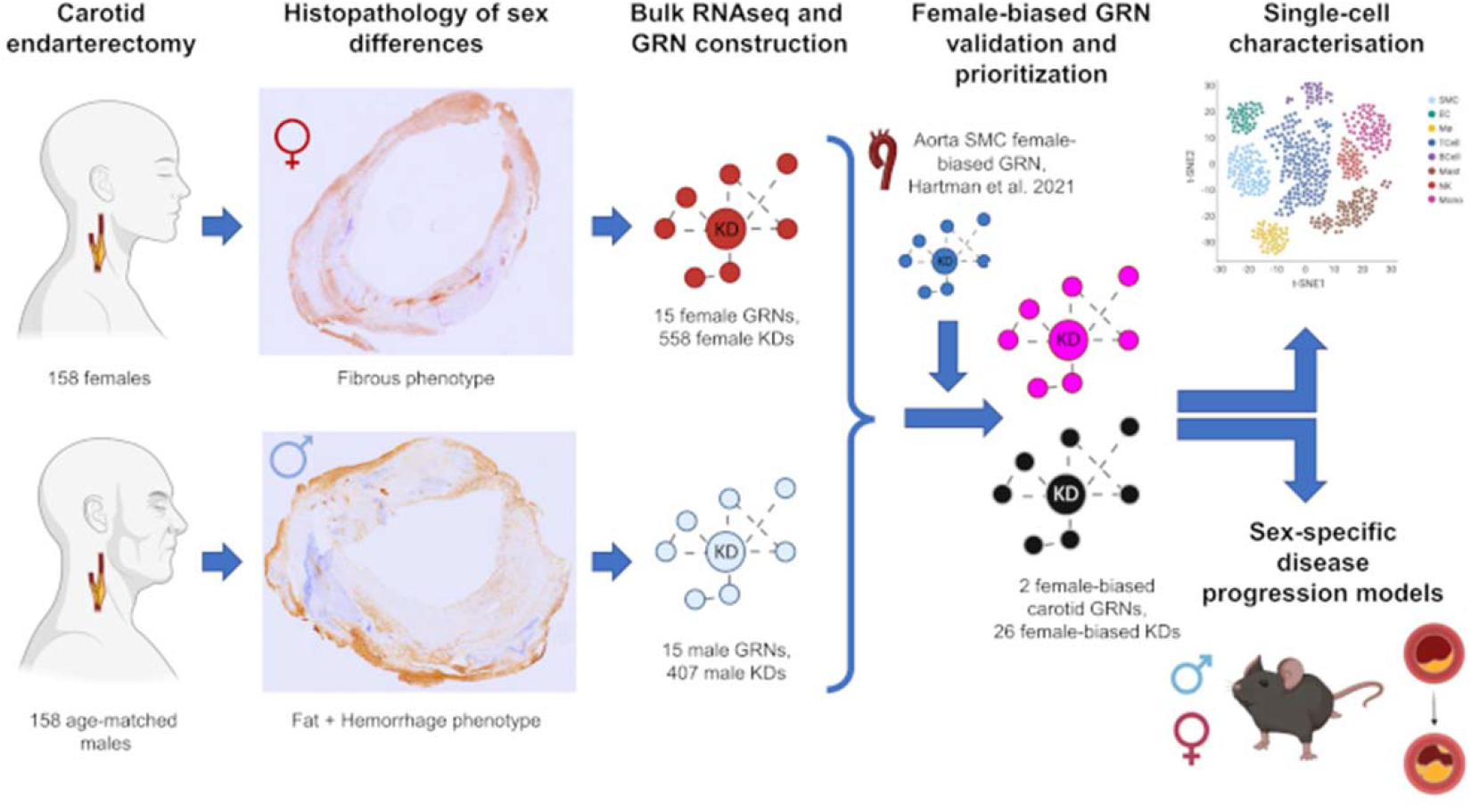
Central illustration of the study. Cartoon images created using BioRender.

### Construction and prioritization of female GRNs in carotid atherosclerotic plaques

*De novo* unbiased co-expression networks were constructed in a sex-stratified manner using gene expression data from advanced plaques which revealed 15 modules in the male group and 15 modules in the female group (**Figure S2, Table S2 and S3**). The median module size was 139 genes [Inter-quartile range (IQR): 80 - 1595] in females and 149 genes [IQR: 65.5 - 1704.8] in males. Enrichment for the top biological processes of the female and male modules is presented in **Figure S3 and Tables S4 and S5**. A heatmap representation of the female-defined modules can be seen in **Figure S4A**. In order to identify genes that drive expression of the gene regulatory networks, we used Bayesian Network Inference and determined key driver (KD) genes as previously described^10^. A total of 558 KD genes were identified in female networks and 407 KD genes were identified in male networks at FDR <0.1 (**Table S2 and S3**).

Next, we prioritized our novel late-stage atherosclerotic GRNs in order to identify the female-biased networks that capture disease relevant SMC biology, based on the following:

1. Female-bias in network activity - measured as a significant difference in average connectivity (a proxy for network activity) of the module in females and males separately, 2. Significant gene overlap with the previously identified SMC female-biased GRN^10^ from aorta, and 3. Co-localization of GRNs genes with known CAD genome-wide association loci^15,16^. Seven out of 15 of the female-defined GRNs (47%) presented female-bias, measured as a significantly higher connectivity to the same networks in males (**Figure S4B-G, Table S6**). The gene composition of 3 of the 7 female-biased networks had significant overlap with the previously reported smooth muscle cell female-biased GRN in aortic tissue^10^ (*p*<0.04, **Figure S5A, Table S7**). A total of 409 genes of the 775 genes included in the aortic GRN (46%) were represented in these 3 novel carotid networks, including 38/63 of the previously identified female-specific KD genes^10^ (**Table S2**). The 3 overlapping GRNs were enriched for genes involved in extracellular matrix organization, collagen organization and angiogenesis (GRN_MAGENTA_) and genes involved in muscle contraction, muscle system process and extracellular organization for (GRN_BLACK_), GRN_TURQUOISE_ was enriched for RNA processing related processes (all *p* <0.001, **Figure S4H** and **Table S4-5**).

Genome-wide association studies (GWAS) have been performed to elucidate the causal variants underlying the mechanisms of both Mendelian and complex diseases. As part of this effort several loci associated with CAD have been identified in previous GWAS studies^15,16^. Therefore, we investigated the enrichment of GRNs with genes within loci associated with CAD^15,16^. Two out of the three female-biased networks (GRN_MAGENTA_ and GRN_BLACK_) were enriched for genes within CAD loci (*p* <0.001, **Table S8 and Figure S5B**). Using this prioritization approach, GRN_MAGENTA_ and GRN_BLACK_ were prioritized for downstream analysis **(Figure 2 and S3I)**.

**Figure 2.**
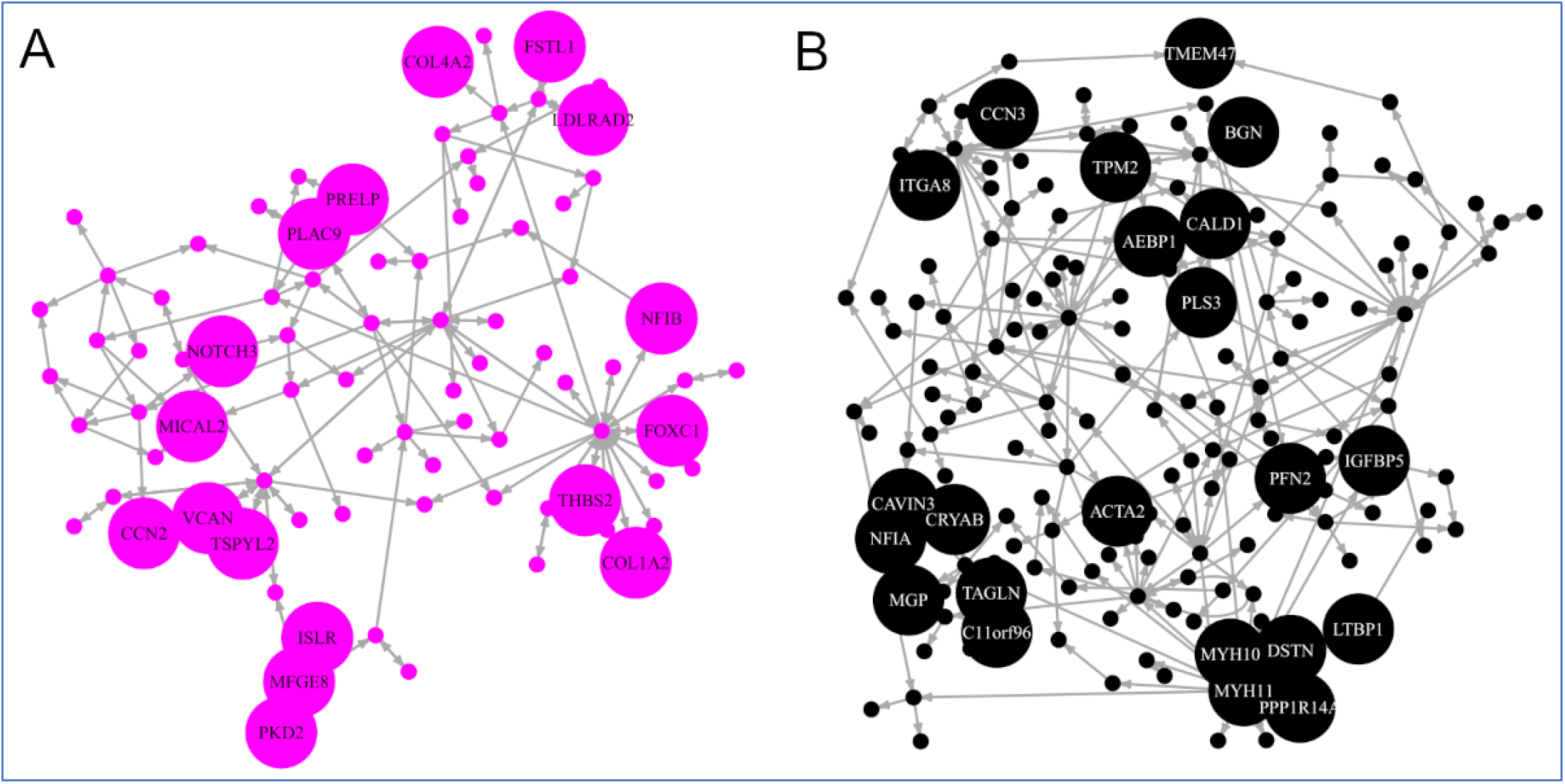
Two prioritized female-biased carotid Gene Regulatory Networks (GRNs) based on overlap with previously identified GRNs and enrichment for CAD GWAS loci. A. GRN_MAGENTA_: 82 genes, 17 KD genes (larger nodes of the network, FDR<0.1). B. GRN_BLACK_: 171 genes, 22 KD genes (larger nodes of the network, FDR<0.1).

### Prioritized female-biased GRNs represent two SMCs phenotypes that modulate plaque stability in females and may play a role in modulating endothelial cells (ECs) transitions

To study these two prioritized female GRNs in more detail, we focused on the cellular origin of the GRNs. We used an extended single-cell RNA sequencing (scRNAseq) data from carotid plaques of 46 patients (20 females and 26 males, **Figure 3A**)^17,18^ to study the cell-type expression of KD genes from the two prioritized female GRNs (GRN_MAGENTA_ and GRN_BLACK_). A total of 20 cell clusters were identified^18^, these were merged down into 10 cell-type clusters. Expression of KD genes from both networks was highest in SMCs, measured through a module score (see Online methods). However, high expression was also identified in both clusters of ECs (**Figure 3B-C**). More specifically, GRN_MAGENTA_ was expressed similarly in both EC clusters (**Figure 3B**) while GRN_BLACK_ was higher expressed in ECs II (**Figure 3C**).

**Figure 3.**
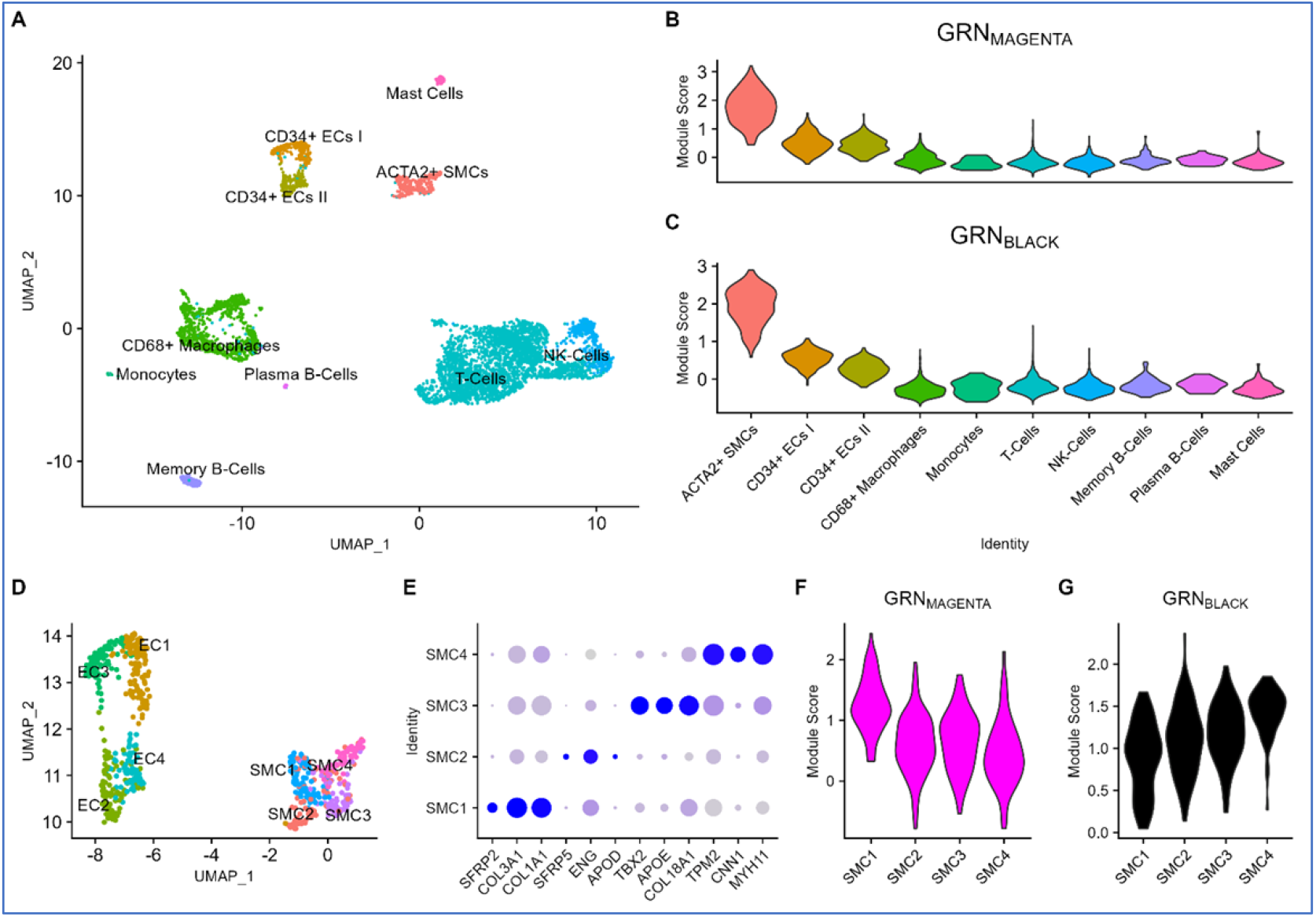
Female-biased Gene Regulatory Networks (GRNs) are expressed in different phenotypes of smooth muscle cells (SMCs) including a myofibroblast-like phenotype: A. UMAP plot of 4,948 single cells from carotid plaque (46 patients, 26 males and 20 females). B. Module score (see Online methods) expression of GRN_MAGENTA_ KD genes in carotid plaque cell-types. C. Module score expression of GRN_BLACK_ KD genes in carotid plaque cell-types. D. Zoom-in UMAP plot of 672 smooth muscle and endothelial single cells and their clusters from carotid plaque (46 patients, 26 males and 20 females). E. Top 3 DEGs for each SMC sub-type, circle size represents percentage of cells expressing the gene and blue intensity represents the normalized expression level. F. Module score expression of GRN_MAGENTA_ KD genes in SMC sub-types. G. Module score expression of GRN_BLACK_ KD genes in SMC sub-types.

Next, we examined the sub-cell-type clusters in SMCs (**Figure 3D-E, Table S9)**. GRN_MAGENTA_ was higher expressed in *KLF4*^+^ SMC1 sub-type compared to the other SMC subtypes (**Figures 3F and S4**). This SMC subtype matches previously reported descriptions of phenotypically modulated myofibroblasts^19^ expressing both classic SMCs markers *ACTA2, TAGLN* and *MYL9*, and fibroblast markers such as *FBLN1, DCN* and *SFRP2* (**Figure S7, Table S9**). Expression of GRN_MAGENTA_ by myofibroblast-like cells was also supported in additional scRNAseq available datasets from carotid endarterectomy^20^ and from coronary transplant patients^21^. Expression of the KD genes from GRN_MAGENTA_ was highest in phenotypically modulated fibrochondrocytes in the carotid dataset^20^ (**Figure S8A-B**) and in modulated TNFRSF11B^+^ fibromyocytes (**Figure S9**) in the coronary tissue^21^ (**Figure S8D-E**). Furthermore, sex-typing of the cells in these datasets using sex-chromosome gene expression revealed that the proportion of *TNFRSF11B*^*+*^ cells over other SMC-like cells was higher in females than in males in both cases (**Figures S10 and S11**). In contrast, GRN_BLACK_ was higher expressed in *TPM2*^+^*CNN1*^+^*MYH11*^+^ SMC4 contractile subtype compared to the other clusters of SMC in the Athero-Express carotid cells, with SMC1 presenting the lowest expression (**Figure 3G**). GRN_BLACK_ KD gene expression was in both cases found highest in canonical SMCs clusters from the complementary scRNAseq datasets (**Figures S8C, S8F, S9B and S9D**).

Modulated SMCs have been suggested to play an atheroprotective role in plaque progression^21^, however they also give rise to other detrimental cellular states, such as osteogenic SMCs^12,22–24^ and foam cells^25–28^. In order to elucidate the role of the two GRNs in shaping plaque phenotypes in females, we assessed the plaque histological characteristics of two groups of plaques based on the dominant expression of either of the networks. In females, 82 out of 158 (52%) plaques had higher expression of GRN_MAGENTA_ compared to GRN_BLACK_ (**Figure S12**). These female patients presented with a more vulnerable and atheromatous plaque (Plaque Vulnerability Index: 3.4 vs 2.8, p <0.001; Atheromatous features: 70% vs 45%, p = 0.005) and a non-significant trend towards more intraplaque haemorrhage present (IPH present: 60% vs 44%, p = 0.067), less ACTA2+ SMCs content (rank-normalized SMC content = 0.06 vs. 0.37, p = 0.074) and more severe symptoms (Stroke symptoms: 33% vs 19%, p = 0.074) (**Table 1**).

**Table 1.** Clinical and carotid plaque characteristics of 158 female patients with different expression of smooth muscle cell (SMC)-related Gene Regulatory Networks (GRNs).

| Characteristic | GRN<sub>MAGENTA</sub> ><br>GRN<sub>BLACK</sub><br>(N = 82) | GRN<sub>BLACK</sub> ><br>GRN<sub>MAGENTA</sub><br>(N = 76) | P-value |
| --- | --- | --- | --- |
| Age (years; mean (SD)) | 69 (10.7) | 67.1 (8.3) | 0.220 |
| Smoking = yes (%) | 31 (39.7) | 38 (50.7) | 0.232 |
| BMI (kg/m<sup>2</sup>; median [IQR]) | 26.1 [23.9, 28.2] | 26.8 [24.2, 29.6] | 0.809 |
| Hypertension = yes (%) | 69 (84.1) | 64 (84.2) | 1.000 |
| Diabetes Mellitus = yes (%) | 12 (14.6) | 17 (22.4) | 0.294 |
| GFR (mL/min; mean (SD)) | 71.3 (20.5) | 73.6 (21.8) | 0.497 |
| Systolic BP (mmHg; mean (SD)) | 152.5 (25) | 156.5 (23.8) | 0.335 |
| Diastolic BP (mmHg; mean (SD)) | 80.6 (12.3) | 83.5 (13) | 0.197 |
| Statin use = yes (%) | 62 (76.5) | 56 (73.7) | 0.818 |
| Stroke symptoms = yes (%) | 26 (32.5) | 14 (18.7) | 0.075 |
| Symptoms (%) |  |  | 0.067 |
| Asymptomatic | 9 (11.2) | 11 (14.7) |  |
| Ocular | 11 (13.8) | 21 (28.0) |  |
| TIA | 34 (42.5) | 29 (38.7) |  |
| Stroke | 26 (32.5) | 14 (18.7) |  |
| Plaque phenotype (%) |  |  | **0.005** |
| Fibrous | 24 (30.0) | 41 (54.7) |  |
| Fibroatheromatous | 36 (45.0) | 25 (33.3) |  |
| Atheromatous | 20 (25.0) | 9 (12.0) |  |
| Plaque vulnerability index (mean (SD)) | 3.4 (1.1) | 2.8 (1.2) | **<0.001** |
| Plaque Hemorrhage = yes (%) | 48 (60.0) | 33 (44.0) | 0.067 |
| SMC content rank-normalized (mean (SD)) | 0.06 (1.05) | 0.37 (1.10) | 0.074 |
| MΦ content rank-normalized (mean (SD)) | 0.11 (1.08) | -0.15 (1.09) | 0.131 |
| High-sensitivity CRP (mg/L, median [IQR]) | 3.3 [1.7, 9.2] | 3 [1.3, 6] | 0.371 |
| HDL (mmol/L; median [IQR]) | 1.2 [0.9, 1.3] | 1.2 [1, 1.5] | 0.520 |
| LDL (mmol/L; mean (SD)) | 2.70 (1.01) | 2.88 (1.17) | 0.470 |

SMC= Smooth Muscle Cells, MΦ= Macrophages, HDL= High-density lipoprotein, LDL= Low-density lipoprotein, CRP= C-reactive protein, TIA= Transient Ischemic Attack, GFR= Glomerular Filtration Rate, BMI= Body Mass Index, BP= Blood Pressure

### Female-biased GRNs are expressed in ECs with signs of EndMT

We previously identified EndMT as a main enrichment in the female-biased SMC gene regulatory network^10^. This suggests a potential role of EndMT in driving sex-differences in atherosclerosis. We therefore investigated the expression levels of our candidate GRNs in ECs sub-types (**Figure 3D**). GRN_MAGENTA_ and GRN_BLACK_ expression levels were highest in EC1 compared to other EC sub-types which represents a *SULF1*^+^ (**Figure 5A**) subtype (of cells that are *CD34*^+^) and are positive for *ACTA2*^+^ (**Figure S6**). EC1 also counts *EFEMP1, IGFBP3* and *DCN* (**Figures 4B and 4C, Table S9**) amongst their top differentially expressed genes, which are fibroblast-like markers. In addition, gene ontology enrichment for this specific subtype of ECs identified enrichment for *TGFβ* response, SMC proliferation and endothelial to mesenchymal transition (*p* <0.001 for GO term enrichment, **Table S11**) further suggesting EndMT may be a player in sex-differences in atherosclerosis.

**Figure 4.**
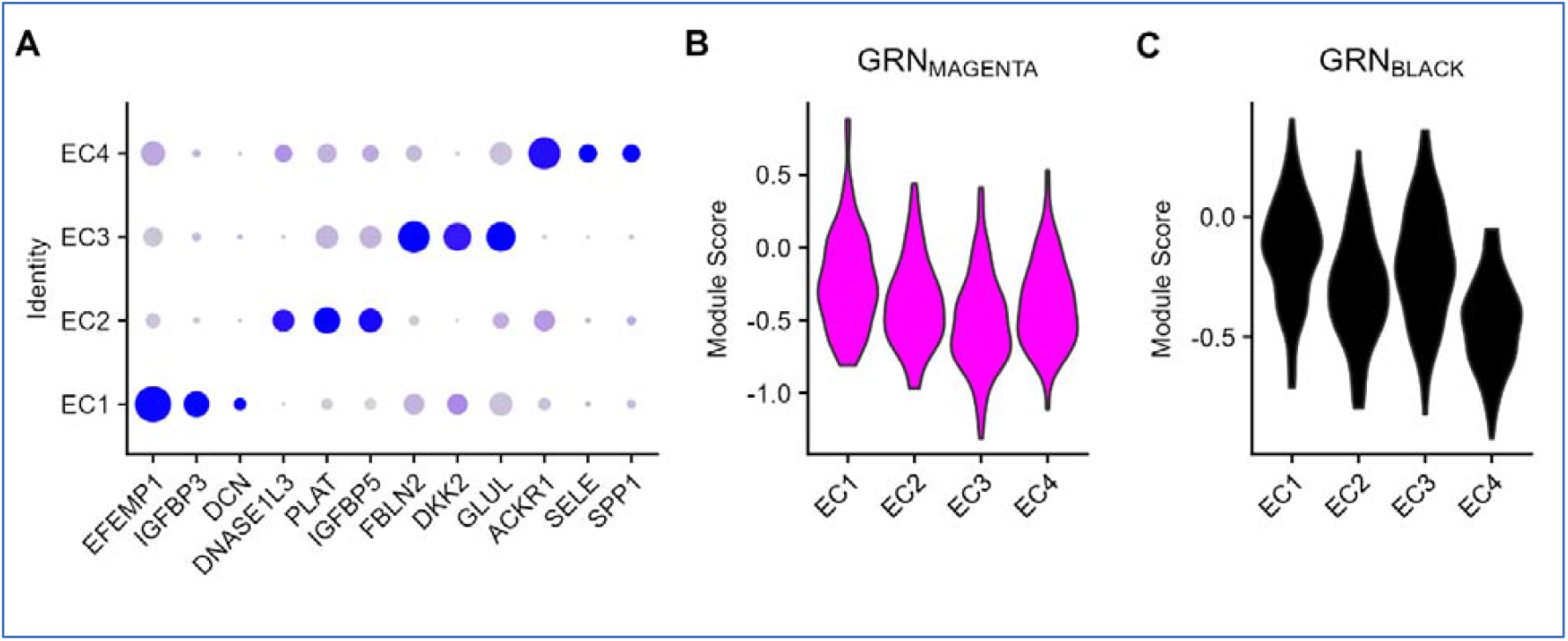
Female-biased Gene Regulatory Networks (GRNs) are expressed in endothelial cells (ECs) with strong signals of endothelial to mesenchymal transition: A. Top 3 differential expressed genes for each EC sub-type, circle size represents percentage of cells expressing the gene and blue intensity represents the normalized expression level. B. Module score expression of GRN_MAGENTA_ KD genes in EC sub-types. C. Module score expression of GRN_BLACK_ KD genes in EC sub-types

**Figure 5.**
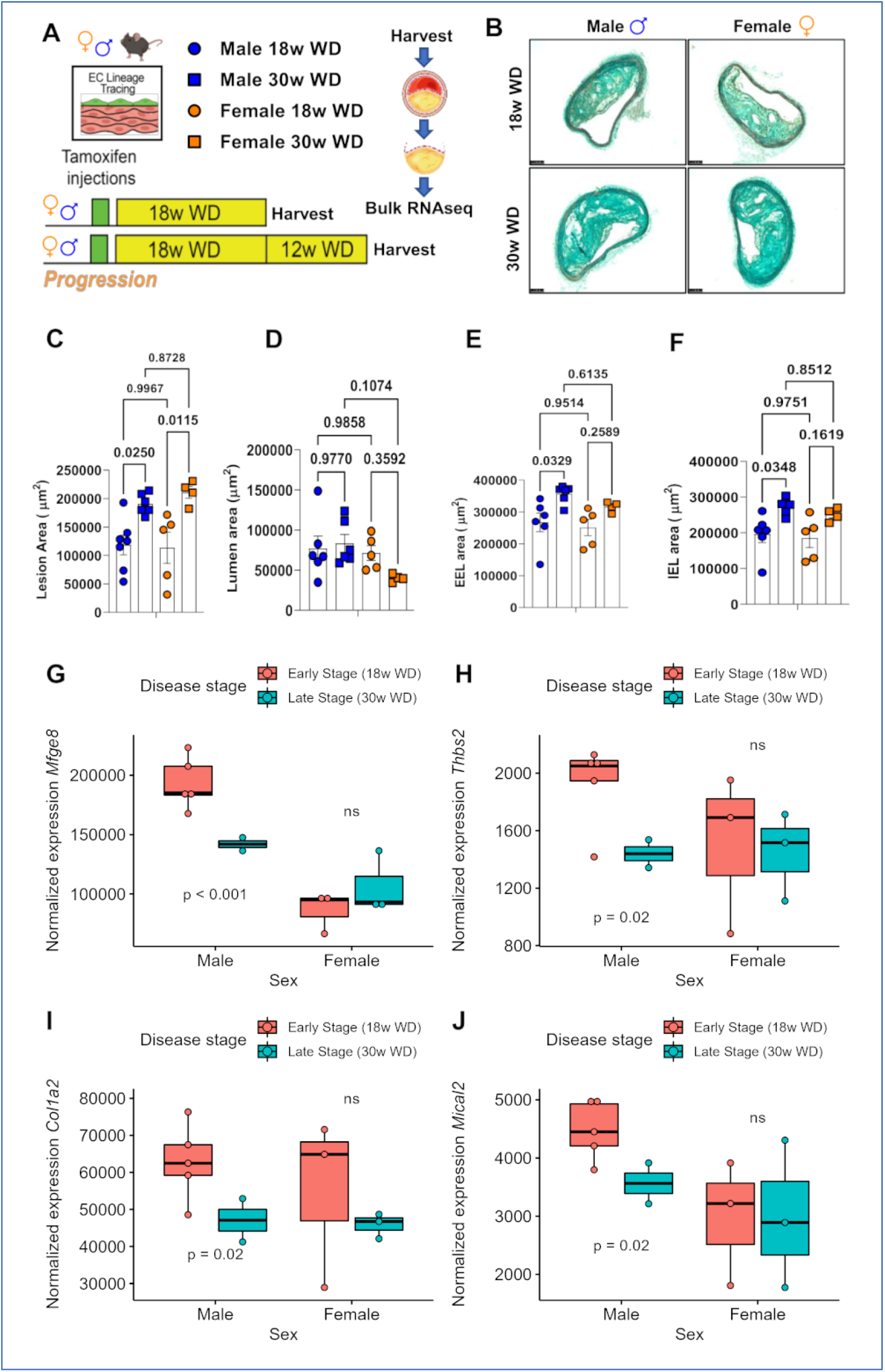
Male-specific downregulation of key driver genes from GRN_MAGENTA_ is observed during atherosclerosis progression in brachiocephalic artery (BCA) lesions from Apoe-/-mice fed a western diet. A. Diagram for experimental design in both male and female mice fed western diet (WD) for 18 weeks (w) and 30w. B. MOVAT staining of representative lesions for male and female mice at 18w and 30w of WD. C. Lession area of male and female mice at 18w WD and 30w WD. D. Lumen area of male and female mice at 18w WD and 30w WD. E. External elastica lamina (EEL) area of male and female mice at 18w WD and 30w WD. F. Internal elastica lamina (IEL) area of male and female mice at 18w WD and 30w WD. G. Normalized expression of *Mfge8* gene by sex and disease stage. H. Normalized expression of *Thbs2* gene by sex and disease stage. C. Normalized expression of *Col1a2* gene by sex and disease stage. D. Normalized expression of *Mical2* gene by sex and disease stage.

### Expression of key driver female-biased genes is maintained in female Apoe^-/-^ mice during disease progression but downregulated in males

To study the role of our prioritized female GRNs in atherosclerosis progression, we used female and male Apoe^-/-^ mice (n = 9 and 13, respectively). Mice were fed a western diet for 18 weeks (18w western diet, i.e., early-stage atherosclerosis) or 30 weeks (30w western diet, i.e., late-stage atherosclerosis, **Figure 5A)**. Lesion size was assessed using Movat staining. A significant increase in lesion size was observed both in males and females in late stages when compared to early atherosclerosis stages (*p* <0.02, **Figure 5B-C**) and no significant difference was observed in lumen size (**Figure 5D**). Brachiocephalic arteries were successfully harvested from a total of 17 mice from an independent experiment (female = 9, male = 8), and the lesion site was successfully laser micro-dissected and processed using bulk RNAseq (female n= 6, male n= 7, see Online methods; **Figure 5A**). A total of 13,083 transcripts out of 21,116 mice transcripts recovered overlapped with the human genes used in our carotid study, including 75/82 (91%) and 160/171 (94%) of GRN_MAGENTA_ and GRN_BLACK_ genes respectively. We performed sex-stratified differential gene expression analysis (DGEA) between conditions (late-stage vs early-stage atherosclerosis) and DGEA between sexes (Male vs Female) at both disease stages (**Tables S11-15**). Enrichment analysis for the differentially expressed genes (DEGs) from the atherosclerotic lesions in male mice identified a male-specific upregulation of calcification and osteogenic processes, translation and RNA processing in late stages and downregulation of ECM organization genes (**Figure S13, Table S17**). Amongst these, expression of KD genes of GRN_MAGENTA_ (including *Col1a2, Thbs2, Mical2* and *Mfge8)* was significantly downregulated in late-stage atherosclerosis in male mice, while expression was maintained in female plaques (**Figure 5G-J**). These results suggest a male-specific decrease in the expression of GRN_MAGENTA_ KD genes during disease progression. In contrast, these genes’ expression is maintained in female plaque potentially shaping the observed sex differences in plaque phenotypes.

## Discussion

Here, we demonstrate that two female GRNs that are both highly expressed in SMCs and ECs, associate with plaque vulnerability and composition, and remain stable in expression during atherosclerosis progression in female mice, and not in male mice. Our data show that fibrous female atherosclerosis is characterized by phenotypic modulation of SMCs and driven by GRNs that promote a vulnerable fibrous plaque rich in extracellular matrix-producing myofibroblast-like SMCs. We highlight the importance of sex-stratification in atherosclerosis research to find new mechanisms of disease progression.

At the gene expression level, we recently uncovered sex-difference in gene connectivity in atherosclerotic tissue which pointed to SMC and ECs as main players in female atherosclerosis^10^. In this study, we substantiate that this dominant female SMC and EC biology is not only important for arterial tissue from coronary artery disease patients but also for carotid plaques from stroke patients. Our mouse data supports the idea that there are intrinsic differences in male and female atherosclerosis pathology progression, as confounding by different risk factor distributions does not play a role in our in-vivo experimental setting. As sex hormones and chromosomes determine biological sex, it may be that the XX sex chromosome complement accelerates atherosclerosis in females, as was found in the murine four core genotype model^29^. Besides the potential role of the sex chromosome complement, estrogen receptor signaling was also previously found in enrichment analyses of female atherosclerotic plaques of women of an average age of 70 years, long after menopause^10^. This may imply that estrogen may still be an important factor to consider in post-menopausal women in vascular tissues, and suggests long-term epigenetic effects.

In line with our previous work, female-biased GRNs in carotid plaques pointed to SMC biology. However, we now have evidence that the previously identified female SMC GRN^10^ might encompass larger diversity at late stages of atherosclerosis development, with two main networks identified in symptomatic atherosclerotic plaques. GRN_MAGENTA_ overlaps in expression with modulated SMCs and points towards a more vulnerable plaque phenotype including higher prevalence of atheroma, which reflects the dual role that modulated SMCs might play in lesion stability depending on their end-stage fate. It is worth noting other factors, such as the higher inflammation observed in male plaques compared to females, as well as known differences in vessel size, and therefore may also shape the fate of SMCs, and potentially ECs also, in the vessel wall.

In this study, we collected plaque samples in the last stage of atherosclerosis which represents the sum of accumulated disease progression over time. While sex differences in atherosclerotic plaques were described to be especially pronounced at younger ages (<50 years) ^30^, the female plaques in our cohort were consistently more fibrous compared to male plaques^3,8^.

In-depth characterization of the GRNs co-expression patterns in scRNAseq from carotid plaques highlighted the relevance of the two most biased networks in females as important for different SMC subtypes or sub-phenotypes. GRN_MAGENTA_ is highly expressed in a myofibroblast-like *KLF4*^+^ SMC subtype and GRN_BLACK_ is mainly expressed in contractile SMCs. In line with this, previous studies have identified female KD genes such as *FN1* to be modulated by *KLF4*^10^ which is the main transcription factor modulating synthetic SMCs, an intermediate state that precedes the myofibroblast phenotype in several models^24^. Our results further prove the importance of SMC plasticity in female atherosclerosis.

We have also identified that KDs of the GRN_MAGENTA_ are highly expressed in a subtype of ECs with biological signatures pointing to SMC proliferation and EndMT. This combined with the high expression of *ACTA2*^+^ in this same subtype suggests that this network may also play a role in EndMT and may contribute to plaque formation in females. However, studying the exact contribution of the SMCs-driven and EC-driven network expression and their potential interactions would require more in-depth research. We postulate that advanced experimental settings such as scRNAseq data from lineage-traced mice in which the origin of the cells expressing these networks can be determined are an important next step to understand if mechanisms that lead to these modulated cell-types are sex-specific.

It is worth noting that ECM proteins which are mainstays of both female-biased carotid GRNs highlighted in this study have been identified as key players within other disease mechanisms that present a marked sex-bias, such as heart failure with preserved ejection fraction (HFpEF)^31^ and myocardial remodelling^32^. Furthermore, several of these genes have been shown to affect SMC phenotypic switching specifically, such as *FN1*^33^ which is also a validated CAD GWAS gene^34^. The key driver *MFGE8* which is a CAD GWAS gene^35^, have been suggested to interact with elastin^36^ and a common loss-of-function variant in a finish population has been shown to associate with atheroprotection^37^. Therefore, the importance of these ECM genes as mediators of sex-specific disease mechanisms should be further studied. The results of our study suggest that while ECM production (including several GRN_MAGENTA_ KD genes) is maintained in female mice’s atherosclerosis progression, its production is actually downregulated in male mice. One hypothesis is that cell-types (i.e. myofibroblasts) producing these ECM proteins are less prevalent in male mice plaques. Alternatively, the networks themselves could be less active in male cells as atherosclerosis progresses. It is also worth noting, that while ECM-producing myofibroblasts are critical in the formation of the fibrous cap, in this study they were found associated with vulnerable plaques in females, suggesting the role of these ECM-producing myofibroblasts in advanced-stages might be detrimental, for example by becoming a scaffold for calcification^38^. In order to answer these questions, one would require high-quality male and female plaque scRNAseq data in both human and mice, preferably within the Four Core Genotypes mouse model to disentangle chromosome from gonadal sex^29^.

This study had some limitations, the biobank we used presents a predominance of male samples (30% females) which resulted in down-sampling of the male cohort to have similar powered studies in females and males, this reduced the overall power of our studies. The use of semi-quantitative phenotypes measured in histological sections, despite having good replication metrics^13,39,40^ may be considered a limitation given the resolution in comparison to the highly complex gene expression patterns in this study. In addition, there is no vascular control tissue in the Athero-Express biobank. However, for the overlapping female-biased GRNs studied here, relevance for disease compared to healthy tissue was assessed in a previous study^10^ using non-atherosclerotic mammary arteries from the same patients, so a degree of relevance for diseased tissue will be applicable here given the overlap found for the networks. We also acknowledge the fact that tamoxifen, which is used in the EC-lineage tracing mice is an estrogen receptor modulator that has been shown to increase cardiovascular risk in women under long-term treatment for breast cancer^41^, however the dose and duration of treatment (a daily injection of 1 mg during 10 days at the age of 6 weeks) is thought not to have a meaningful impact on atherosclerotic disease development over the next 30 weeks as effects of tamoxifen are observed over periods of years in humans^41^. Furthermore, in this study mice used were 6 weeks at the start of the experiment, and while our human studies are performed on advanced lesions, the atherosclerosis observed in mice is progressing and shows a similar trajectory of differences between males and females as observed in humans (i.e. higher ECM in females than males).

In conclusion, this study brings us closer to understanding the mechanisms that lead to the observable differences between male and female atherosclerotic plaques, as well as sheds light into how these mechanisms might be regulated during atherosclerosis disease progression. We shed light on possible new gene targets involved in SMC plasticity in female atherosclerosis and provide the foundation to identify targets for novel therapies which might inhibit detrimental SMC transitions and further stabilize the female plaque. Furthermore, in the context of the current interest in molecular diagnostics, our results might help identify sex-specific targets which might prove useful to close the gap in cardiovascular disease diagnosis between males and females and bring the promise of personalized medicine a step closer.

## Online Methods

### Human patient samples

Patients undergoing endarterectomy of the carotid artery in two Dutch tertiary referral centers between 2002 and 2020 were included in this study. Study procedures comprise of a baseline blood withdrawal, an extensive questionnaire filled in by the participants verified against medical records, and collection of carotid arterial plaque material during surgery. All patients provided written informed consent before surgery, the study was approved by the Local Medical Ethical Committee and conducted according to the Declaration of Helsinki^42^. Details of the Athero-Express study protocol have been described previously^13,40,43^. All available plaque bulk RNA female samples were used in this study, male samples were down-sampled by splitting the male samples in 10-year groups and selecting random males in each group to match the number of females in the same age groups.

### Mice

The University of Virginia Animal Care and Use Committee approved animal protocols (Protocol 2400). The Cdh5-Cre ERT2 R26R-eYFP Apoe-/-mice (Species: *Mus musculus*, Age: 6 weeks at the time of tamoxifen treatment, Sex: 13 males and 9 females, Strain: C57BL/6J, Source: Jackson laboratory) used in the present study have been described in our recent study^44^. The animals were randomized after genotyping at the age of 6 weeks before starting the experiment. All experiments were performed blindly. Cdh5 Cre ERT2 R26R-eYFP Apoe-/-mice, Cre recombinase was activated with a series of ten tamoxifen injections (1mg/day/mouse; Sigma Aldrich, T-5648) over a 2-week period. One week after the tamoxifen treatment, mice were switched from a normal chow diet (Harlan Teklad TD.7012) to a high fat Western-type diet (WD), containing 21% milk fat and 0.15% cholesterol (Harlan Teklad; TD.88137) for 18 weeks or 30 weeks.

### Atherosclerotic plaque morphometry of mouse BCA lesions

Paraformaldehyde-fixed paraffin-embedded BCAs were serially cut into 10 μm thick sections from the aortic arch to the bifurcation of the right subclavian artery. For morphometric analysis, we performed modified Russell-Movat staining on the BCA at 750 μm. The lesion, lumen, external elastic lamina (outward remodeling), and the internal elastic lamina area were measured on digitized images of the Movat staining using Fiji software version 1.53c. For lesion area, lumen area, EEL area, and IEL area one-way ANOVA method in GraphPad Prism 9.4.1 version was used.

### Human plaque histology

As described previously^13,39,40^, the atherosclerotic plaque was processed directly after surgery and (immune-)histochemical staining was routinely performed on the culprit lesion (segment with the highest plaque burden) for identification of macrophages (CD 68), calcification (haematoxylin-eosin (HE)), smooth muscle cells (alfa actin), collagen (Picro Sirius red (PSR)), plaque hemorrhage (HE, Elastin von Gieson staining), vessel density (CD34) and fat (PSR, HE). Assessment of overall plaque vulnerability was performed as previously described by Verhoeven et al.^13^. Briefly, macrophages and smooth muscle cells were semi-quantitatively defined as no/minor or moderate/heavy. Each plaque characteristic that defines a stable plaque (i.e., no/minor macrophages, moderate/heavy collagen, moderate/heavy smooth muscle cells and <10% fat) was given a score of 0, while each plaque characteristic that defines a vulnerable plaque (i.e., moderate/heavy macrophages, no/minor collagen, no/minor smooth muscle cells and ≥10% fat) was given a score of 1. The score of each plaque characteristic was summed resulting in a final plaque score ranging from 0 (most stable plaque) to 4 (most vulnerable plaque). Intraobserver and interobserver variability were examined previously and showed good concordance (κ = 0.6–0.9)^45^.

### Bulk RNA sequencing of human carotid plaques

As the culprit lesion is used for plaque histology following the standardized Athero-Express protocol^13^, the adjacent plaque segments were used for RNA sequencing. To measure bulk RNA expression in the plaques total RNA was isolated according to the manufacturers protocol after processing of the plaque segments using ceramic beads and tissue homogenizer (Precellys, Bertin intruments, Montigny-le-Bretonneux) with use of TriPure (Sigma Aldrich). After precipitating RNA in the aqueous phase with propanolol, RNA was washed with 75% ethanol and either used immediately after an additional washing step with 75% ethanol or stored in 75% ethanol for later use. Subsequently, library preparation was performed as described before^46–48^. Ethanol was removed and the pellet air-dried. Then, primer mix (5ng primer per reaction) was added to initiate primer annealing at 65 degrees Celsius for 5 minutes. Subsequently, reverse transcription (RT) was executed. Subsequent reverse-transcription RT reaction; first strand reaction for 1hour at 42 °C, heat inactivated for 10 minutes at 70°C, second strand reaction for 2 hours at 16 °C, and then put on ice until proceeding to sample pooling.

This initial RT reaction used the following primer design: an anchored polyT, a unique 6bp barcode, a unique molecular identifier (UMI) of 6bp, the 5’ Illumina adapter and a T7 promoter, as described (57). Each sample now contained its own unique barcode making it possible to pool together cDNA samples at 7 samples per pool. Complementary DNA (cDNA) was cleaned using AMPure XP beads (Beckman Coulter), washed with 80% ethanol, and resuspended in water before proceeding to the in vitro transcription (IVT) reaction (AM1334; Thermo-Fisher) incubated at 37 °C for 13 hours. Next, Exo-SAP (Affymetrix, Thermo-Fisher) was used to remove primers, upon which amplified RNA (aRNA) was fragmented, cleaned with RNAClean XP (Beckman-Coulter), washed with 70% ethanol, air-dried, and resuspended in water. RNA yield and quality in the suspension were checked by Bioanalyzer (Agilent) after removal of the beads with use of a magnetic stand. By performing an RT reaction using SuperScript II reverse transcriptase (Invitrogen/Thermo-Fisher) according to the protocol of the manufacturer cDNA library construction was initiated. Next, PCR amplification was performed as described previously ^46–48^. PCR products were cleaned twice using AMPure XP beads (Beckman Coulter). Qubit fluorometric quantification (Thermo-Fisher) and Bioanalyzer (Agilent) were used to checked Library cDNA yield and quality. Illumina Nextseq500 platform was used to sequence the libraries; paired end, 2 × 75bp. After sequencing, retrieved fastq files were de-barcoded and split into forward and reverse reads. From there, the reads were mapped using Burrows-Wheel aligner (BWA37) version 0.7.17-r1188, calling ‘bwa aln’ with settings -B 6 -q 0 -n 0.00 -k 2 -l 200 -t 6 for R1 and -B 0 -q 0 -n 0.04 -k 2 -l 200 -t 6 for R2, ‘bwa sampe’ with settings -n 100 - N 100, and a cDNA reference (assembly hg19, Ensembl release 84). Read and UMI counts were acquired from SAM files with use of custom perl code and collected into count matrices. Further analyses were performed using R^49^ version 3.6.2 and later and its IDE Rstudio^50^ version 1.2 and later. Genes were annotated with Ensembl ID’s, basic quality control was performed (filtering out samples with low gene numbers (<10,000 genes) and read counts (<18,000 reads)).

### Gene regulatory networks analysis in human

The software package WGCNA^51^ (v. 1.69) was used to generate modules of co-expression genes on the 486 available men RNAseq samples. After excluding all the ribosomal genes and included only the protein-coding genes with annotated HGCN names, a set of 12,765 protein-coding genes which passed quality control (average >1 count per sample) was used for module generation. The raw read counts were corrected for UMI sampling:

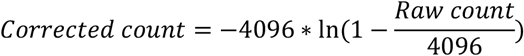

then normalized by sample sequencing depth and log-transformed. A signed network was constructed using the robust “bicor” correlation measure. To determine the exponent used for the adjacency matrix construction, soft thresholding analysis was performed with the WGCNA package for powers ranging from 2 to 30. The cut-off for assuming scale-free topology was set at an R-squared of 0.8, while having a median connectivity of lower than 100. The chosen lowest complying power was 24 (**Figure S1**). The network was constructed by first generating an adjacency matrix, which was transformed into a topological overlay matrix^51^. Modules were detected by clustering the average distance of the dissimilarity matrix (defined as 1-topological overlay matrix) and cutting the subsequent dendrogram by using the *cutreeDynamic* function (*deepSplit =3, minClusterSize = 20*). Modules were named after colors as provided by the WGCNA package, for each analysis Gray module was excluded as it is considered a “bin” module where genes that cant be assigned to a module are placed. Module eigengenes were calculated by taking the first principal component of gene expression in that module. The module eigengenes were correlated to clinical traits by Pearson correlation, with a Student asymptotic *p*-value test for significance.

### Bayesian inference in gene regulatory network and key driver analysis

Bayesian network inference was performed as described previously^10^, in brief, the rcausal package (v0.99.0).The Fast Greedy Equivalence Search (max degrees = 100) for continuous data algorithm was used to generate the Bayesian network on the WGCNA-generated modules. Key driver analysis (KDA) was performed as integrated in the Mergeomics package (v1.10.0), number of permutations was set to 20. Key drivers were considered significant if FDR < 0.1.

### Bulk RNAseq of BCA lesion microdissections from Apoe^-/-^ mice

Total RNA was isolated using RNeasy kit (Qiagen 74106) from the micro-dissected BCA lesions of Cdh5 Cre ERT2 R26R-eYFP Apoe-/-mice fed a high-fat Western diet for 18 weeks (18w) or 30 weeks (30w). The RNA library was prepared according to manufacturer instructions with rRNA reduction with an average RNA integrity number (RIN) of 6.6. A total of 17 BCA RNA samples were isolated from mice (male 18w= 5, male 30w = 3, female 18w = 5 and female 30w = 4). A total of 4 samples were discarded due to low yield of RNA libraries, resulting in a total of 13 RNA samples for analysis (male 18w= 5, male 30w = 2, female 18w = 3 and female 30w = 3). RNA libraries were provided to the company Novogene Inc. (USA) for bulk RNAseq. Libraries were sequenced with the Illumina short-read sequencing (Illumina HiSeq v4; 100 bp and 25 million paired-end reads). Raw reads were QCed to exclude reads containing adapters, reads containing ambiguous calls (N) >10% and reads where 50% of bases had a Qscore <5. QCed reads were mapped to the *Mus musculus* mice reference genome using the STAR software^52^. Reads per transcript were extracted from the STAR output and converted to FPKM (Fragments Per Kilobase of transcript sequence per Millions base pairs sequenced). A total of 54,532 transcripts were obtained of which 21,116 transcripts had an average expression >1 read per sample and were used for differential gene expression. Read counts were normalized and differential gene expression between groups was performed using DESeq2 in R^53^. Reactome gene enrichment analysis for genes with significant up- or down-regulation was performed.

### Single-cell RNAseq of human carotid plaques

Single-cell RNAseq was sequnces following previously published protocols^17,18^. In brief, atherosclerotic lesions were collected from 20 female and 26 male patients undergoing a carotid endarterectomy procedure. All pathological tissue was included in the Athero-Express Biobank Study biobank (AE, www.atheroexpress.nl) at the University Medical Centre Utrecht (UMCU)^13^. After sequencing, data were processed in an R 3.5 and 4.0 environment using Seurat version 3.2.2^54^. Mitochondrial genes were excluded and doublets were omitted by gating for unique reads per cell (between 400 and 10 000) and total reads per cell (between 700 and 30 000). Data were corrected for sequencing batches using the function SCTransform. Clusters were created with 30 principal components at resolution 0.8. Multiple iterations of clustering were used to determine optimal clustering parameters. Population identities were assigned by evaluating gene expression per individual cell clusters. Sub-populations of SMCs were determined by isolating the SMC and EC populations from the complete population of cells mentioned above and re-assigned clusters. Clusters were created with 10 principal components at resolution 1.4. New subtypes of SMC and EC identities were assigned by evaluating the DEGs per cell clusters using enrichment analysis in enrichR (v. 3.0)^55^. Module score for the different set of KD genes were generated using the addModuleScore function in Seurat^54^ which estimates a proxy of expression for all the genes in the provided set.

### Data and scripts availability

Data and scripts are available upon reasonable request.

## Acknowledgments

The authors would like to acknowledge all the participants of the Athero-Express biobank for agreeing to being part of the study. The authors would also like to acknowledge the support from the Young Investigator Award program from the PlaqOmics Network of Excellence within the Leducq Foundation.

## Supplemental Material

Tables S1-S17

Figure S1-S13

## References

1. Canto, J. G. et al. Association of Age and Sex With Myocardial Infarction Symptom Presentation and In-Hospital Mortality. JAMA 307, 813–822 (2012).

2. Vaccarino, V. et al. Sex Differences in Mortality After Myocardial Infarction: Evidence for a Sex-Age Interaction. Arch. Intern. Med. 158, 2054–2062 (1998).

3. de Bakker, M. et al. The age- and sex-specific composition of atherosclerotic plaques in vascular surgery patients. Atherosclerosis 310, 1–10 (2020).

4. Burke, A. P., Farb, A., Malcom, G. & Virmani, R. Effect of menopause on plaque morphologic characteristics in coronary atherosclerosis. Am. Heart J. 141, S58–62 (2001).

5. Farb, A. et al. Coronary plaque erosion without rupture into a lipid core. A frequent cause of coronary thrombosis in sudden coronary death. Circulation 93, 1354–1363 (1996).

6. Burke, A. P. et al. Coronary Risk Factors and Plaque Morphology in Men with Coronary Disease Who Died Suddenly. N. Engl. J. Med. 336, 1276–1282 (1997).

7. Yamamoto, E. et al. Clinical and Laboratory Predictors for Plaque Erosion in Patients With Acute Coronary Syndromes. J. Am. Heart Assoc. 8, e012322 (2019).

8. Hellings, W. E. et al. Gender-associated differences in plaque phenotype of patients undergoing carotid endarterectomy. J. Vasc. Surg. 45, 289–296 (2007).

9. Quillard, T., Franck, G., Mawson, T., Folco, E. & Libby, P. Mechanisms of erosion of atherosclerotic plaques. Curr. Opin. Lipidol. 28, (2017).

10. Hartman, R. J. G. et al. Sex-Stratified Gene Regulatory Networks Reveal Female Key Driver Genes of Atherosclerosis Involved in Smooth Muscle Cell Phenotype Switching. Circulation 143, 713–726 (2021).

11. de Jager, S. C. A. et al. Preeclampsia and coronary plaque erosion: Manifestations of endothelial dysfunction resulting in cardiovascular events in women. Eur. J. Pharmacol. 816, 129–137 (2017).

12. Alencar, G. F. et al. Stem Cell Pluripotency Genes Klf4 and Oct4 Regulate Complex SMC Phenotypic Changes Critical in Late-Stage Atherosclerotic Lesion Pathogenesis. Circulation 142, 2045–2059 (2020).

13. Verhoeven, B. A. N. et al. Athero-Express: Differential Atherosclerotic Plaque Expression of mRNA and Protein in Relation to Cardiovascular Events and Patient Characteristics. Rationale and Design. European Journal of Epidemiology vol. 19 (2004).

14. Mokry, M. et al. Transcriptomic-based clustering of advanced atherosclerotic plaques identifies subgroups of plaques with differential underlying biology that associate with clinical presentation. medRxiv 2021.11.25.21266855 (2021) doi:10.1101/2021.11.25.21266855.

15. Erdmann, J., Kessler, T., Munoz Venegas, L. & Schunkert, H. A decade of genome-wide association studies for coronary artery disease: the challenges ahead. Cardiovasc. Res. 114, 1241–1257 (2018).

16. Aragam, K. G. et al. Discovery and systematic characterization of risk variants and genes for coronary artery disease in over a million participants. medRxiv 2021.05.24.21257377 (2021) doi:10.1101/2021.05.24.21257377.

17. Depuydt, M. A. C. et al. Microanatomy of the Human Atherosclerotic Plaque by Single-Cell Transcriptomics. Circ. Res. 127, 1437–1455 (2020).

18. Slenders, L. et al. Intersecting single-cell transcriptomics and genome-wide association studies identifies crucial cell populations and candidate genes for atherosclerosis. Eur. Hear. J. open 2, oeab043 (2022).

19. Hu, Z. et al. Single-Cell Transcriptomic Atlas of Different Human Cardiac Arteries Identifies Cell Types Associated With Vascular Physiology. Arterioscler. Thromb. Vasc. Biol. 41, 1408–1427 (2021).

20. Pan, H. et al. Single-Cell Genomics Reveals a Novel Cell State During Smooth Muscle Cell Phenotypic Switching and Potential Therapeutic Targets for Atherosclerosis in Mouse and Human. Circulation 142, 2060–2075 (2020).

21. Wirka, R. C. et al. Atheroprotective roles of smooth muscle cell phenotypic modulation and the TCF21 disease gene as revealed by single-cell analysis. Nat. Med. 25, 1280–1289 (2019).

22. Alexopoulos, N. & Raggi, P. Calcification in atherosclerosis. Nat. Rev. Cardiol. 6, 681– 688 (2009).

23. Qiao, J.-H., Mishra, V., Fishbein, M. C., Sinha, S. K. & Rajavashisth, T. B. Multinucleated giant cells in atherosclerotic plaques of human carotid arteries: Identification of osteoclast-like cells and their specific proteins in artery wall. Exp. Mol. Pathol. 99, 654–662 (2015).

24. Yap, C., Mieremet, A., de Vries, C. J. M., Micha, D. & de Waard, V. Six Shades of Vascular Smooth Muscle Cells Illuminated by KLF4 (Krüppel-Like Factor 4). Arterioscler. Thromb. Vasc. Biol. 41, 2693–2707 (2021).

25. Chaabane, C., Coen, M. & Bochaton-Piallat, M.-L. Smooth muscle cell phenotypic switch: implications for foam cell formation. Curr. Opin. Lipidol. 25, 374–379 (2014).

26. Bennett, M. R., Sinha, S. & Owens, G. K. Vascular Smooth Muscle Cells in Atherosclerosis. Circ. Res. 118, 692–702 (2016).

27. Allahverdian, S., Chaabane, C., Boukais, K., Francis, G. A. & Bochaton-Piallat, M.-L. Smooth muscle cell fate and plasticity in atherosclerosis. Cardiovasc. Res. 114, 540– 550 (2018).

28. Wang, Y. et al. Smooth Muscle Cells Contribute the Majority of Foam Cells in ApoE (Apolipoprotein E)-Deficient Mouse Atherosclerosis. Arterioscler. Thromb. Vasc. Biol. 39, 876–887 (2019).

29. AlSiraj, Y. et al. XX sex chromosome complement promotes atherosclerosis in mice. Nat. Commun. 10, 2631 (2019).

30. Yahagi, K., Davis, H. R., Arbustini, E. & Virmani, R. Sex differences in coronary artery disease: Pathological observations. Atherosclerosis 239, 260–267 (2015).

31. Tromp, J. et al. Identifying Pathophysiological Mechanisms in Heart Failure With Reduced Versus Preserved Ejection Fraction. J. Am. Coll. Cardiol. 72, 1081–1090 (2018).

32. Piro, M., Della Bona, R., Abbate, A., Biasucci, L. M. & Crea, F. Sex-Related Differences in Myocardial Remodeling. J. Am. Coll. Cardiol. 55, 1057–1065 (2010).

33. Hu, W.-Y. et al. Phenotypic Modulation by Fibronectin Enhances the Angiotensin II– Generating System in Cultured Vascular Smooth Muscle Cells. Arterioscler. Thromb. Vasc. Biol. 20, 1500–1505 (2000).

34. Klarin, D. et al. Genetic analysis in UK Biobank links insulin resistance and transendothelial migration pathways to coronary artery disease. Nat. Genet. 49, 1392–1397 (2017).

35. Nikpay, M. et al. A comprehensive 1000 Genomes–based genome-wide association meta-analysis of coronary artery disease. Nat. Genet. 47, 1121–1130 (2015).

36. Larsson, A. et al. Lactadherin binds to elastin – a starting point for medin amyloid formation? Amyloid 13, 78–85 (2006).

37. Ruotsalainen, S. E. et al. Loss-of-function of MFGE8 and protection against coronary atherosclerosis. medRxiv (2021) doi:10.1101/2021.06.23.21259381.

38. Schinke, T., McKee, M. D. & Karsenty, G. Extracellular matrix calcification: where is the action? Nat. Genet. 21, 150–151 (1999).

39. de Bakker, M. et al. The age- and sex-specific composition of atherosclerotic plaques in vascular surgery patients. Atherosclerosis 310, 1–10 (2020).

40. Van Koeverden, I. D. et al. Testosterone to oestradiol ratio reflects systemic and plaque inflammation and predicts future cardiovascular events in men with severe atherosclerosis. Cardiovasc. Res. 115, 453–462 (2019).

41. Harvey, H. A., Kimura, M. & Hajba, A. Toremifene: An evaluation of its safety profile. The Breast 15, 142–157 (2006).

42. Adopted by the 18th WMA General Assembly, Helsinki, Finland, June 1964; amended by the 29th WMA General Assembly, Tokyo, Japan, October 1975; 35th WMA General Assembly, Venice, Italy, October 1983; 41st WMA General Assembly, H. K., September 1989; 48th WMA General Assembly, Somerset West, Republic of South Africa, October 1996, and the 52nd WMA General Assembly, Edinburgh, Scotland, O. 2000 & A. World Medical Association Declaration of Helsinki. Bull. World Health Organ. 79(4) (2001) doi:10.1111/ddg.13528.

43. Van Der Laan, S. W. et al. Variants in ALOX5, ALOX5AP and LTA4H are not associated with atherosclerotic plaque phenotypes: The Athero-Express Genomics Study. Atherosclerosis 239, 528–538 (2015).

44. Newman, A. A. C. et al. Multiple cell types contribute to the atherosclerotic lesion fibrous cap by PDGFRβ and bioenergetic mechanisms. Nat. Metab. 3, 166–181 (2021).

45. Hellings, W. E. et al. Intraobserver and interobserver variability and spatial differences in histologic examination of carotid endarterectomy specimens. J. Vasc. Surg. 46, 1147–1154 (2007).

46. Hashimshony, T. et al. CEL-Seq2: Sensitive highly-multiplexed single-cell RNA-Seq. Genome Biol. 17, (2016).

47. Hashimshony, T., Wagner, F., Sher, N. & Yanai, I. CEL-Seq: Single-Cell RNA-Seq by Multiplexed Linear Amplification. Cell Rep. 2, 666–673 (2012).

48. Ferraz, M. A. M. M. et al. An oviduct-on-a-chip provides an enhanced in vitro environment for zygote genome reprogramming. Nat. Commun. 9, (2018).

49. R Core Team (2020). R: A language and environment for statistical computing. R Foundation for Statistical Computing, Vienna, Austria. (2020).

50. R Studio Team (2020). RStudio: Integrated Development for R. RStudio, PBC, Boston, MA (2020).

51. Langfelder, P. & Horvath, S. WGCNA: an R package for weighted correlation network analysis. BMC Bioinformatics 9, 559 (2008).

52. Dobin, A. et al. STAR: ultrafast universal RNA-seq aligner. Bioinformatics 29, 15–21 (2013).

53. Love, M. I., Huber, W. & Anders, S. Moderated estimation of fold change and dispersion for RNA-seq data with DESeq2. Genome Biol. 15, (2014).

54. Stuart, T. & Satija, R. Integrative single-cell analysis. Nat. Rev. Genet. 20, 257–272 (2019).

55. Chen, E. Y. et al. Enrichr: interactive and collaborative HTML5 gene list enrichment analysis tool. BMC Bioinformatics 14, 128 (2013).

